# Quantifying Scientific Consensus in Biomedical Hypotheses via LLM-Assisted Literature Screening

**DOI:** 10.64898/2026.04.06.716861

**Authors:** Uiyun Kim, Ohhyeon Kwon, Doheon Lee

## Abstract

Systematic literature reviews are labor-intensive tasks in biomedical research. While Large Language Models (LLMs) using Retrieval-Augmented Generation (RAG) techniques have enhanced information accessibility, the inherent complexity of biological systems—characterized by high context dependency and conflicting data—remains a primary driver of LLM hallucinations. This imposes a structural constraint that limits the precision of evidence synthesis. To address these limitations, we propose an automated framework designed for the exhaustive identification of supporting and contradictory evidence within a target literature set. Rather than relying on a model’s pre-trained knowledge, our system requires the LLM to review each paper individually to determine its alignment with a specific research hypothesis. By evaluating semantic context, the framework captures subtle contradictions that are often overgeneralized by conventional methods. The framework’s performance was validated using the BioNLI task, where it demonstrated high classification accuracy in distinguishing whether evidence supports or contradicts a given hypothesis. Notably, the implementation of an ensemble approach provided superior stability and slightly higher precision compared to individual models. Furthermore, the framework exhibited robust performance across several well-established biological hypotheses, confirming its practical utility and reliability in real-world research. This approach provides a rigorous basis for biomedical discovery by enabling the precise, systematic analysis of biological literature and the robust collection of evidence.

## 1 Introduction

The rapid advancement of generative artificial intelligence (AI) is transforming the processing and analysis of vast amounts of information across various fields. In modern biomedical research, where millions of articles are published annually, the impact of this technology is becoming increasingly crucial to overcome information overload and synthesize complex knowledge [1]. To address these challenges, various methodological approaches have been proposed. For instance, large-scale language models (LLMs) such as **MedGemma**, fine-tuned on extensive domain-specific datasets, have demonstrated remarkable capabilities in sophisticated medical reasoning and professional question-answering by leveraging a specialized vision-language foundation [2]. Concurrently, specialized encoder-based models like **BioBERT** remain fundamental in structured knowledge synthesis. These models are highly effective in **Named Entity Recognition (NER)** and **Relation Extraction (RE)**, enabling the automated identification of biological entities—including genes, proteins, and diseases—and the extraction of their relations from unstructured text [3].

Furthermore, the integration of these extraction techniques into comprehensive platforms like **PubTator 3** has greatly enhanced the accessibility of biomedical insights. By establishing a large-scale database of relations between biological entities across the entire PubMed corpus, PubTator 3 provides researchers with a comprehensive biological foundation [4, 5]. Such resources go beyond simple text mining by enabling advanced visualization and semantic search, thereby significantly accelerating hypothesis generation and experimental validation in the era of big data.

Traditionally, systematic literature reviews have been the essential to evidence-based biomedical research, as they integrate existing knowledge on diseases and validate biological hypotheses [6]. However, the sheer volume of scientific literature has now exceeded the cognitive capacity of human researchers [7]. Manually reviewing thousands of diverse studies to identify conflicting evidence is a labor-intensive process that creates a significant bottleneck in research. Therefore, developing an automated literature analysis framework is no longer merely optional but a necessity to ensure both the speed and precision of scientific discovery.

The integration of Large Language Models (LLMs) and Retrieval-Augmented Generation (RAG) has transformed academic information retrieval by enabling the synthesis of complex findings through natural language queries [8]. Despite this progress, a fundamental challenge remains in the biomedical domain: the extreme context-dependency of biological systems. Unlike static datasets, biological functions and interactions are dynamic, varying significantly across cellular environments, genetic backgrounds, and disease states [9]. When models rely on generalized linguistic patterns rather than these specific nuances, they prone to hallucinations and critical errors [10].

This limitation is particularly evident when handling conflicting data. Biological context operates within a multi-layered hierarchy—from molecular pathways to tissue structures—where a factor’s role is defined by its specific environment [9]. Current LLMs, however, are rooted in probabilistic token prediction, which inherently favors statistical consensus [10]. This “generalization bias” often dismisses rare but pivotal contradictory evidence as statistical noise, thereby undermining the reliability of predictive modeling in evidence synthesis [10].

Furthermore, the standard practice of splitting documents into chunks in RAG systems can lead to the loss of critical context, as relevant information is often separated and not retrieved together. [11]. Without maintaining the integrity of this biological context, models may present conflicting results as equally valid, leading to logical contradictions that compromise the rigor of systematic literature reviews.

To address these limitations, this study proposes an automated evidence analysis framework that performs an independent, instance-level review of each publication within the target literature. Unlike traditional RAG systems that often lose vital experimental context due to arbitrary document chunking [11], our approach ensures that the LLM evaluates the complete narrative of every paper to identify both supporting and contradictory evidence. By requiring the model to reference specific experimental conditions—such as the choice of cell lines or stimulus timing—the framework prevents the bias toward statistical consensus and visualizes fine-grained logical conflicts that are frequently overlooked by conventional summarization methods.

The performance of this framework was rigorously validated through the Biomedical Natural Language Inference (BioNLI) task, demonstrating high accuracy in classifying how evidence aligns with specific hypotheses [12]. In particular, the introduction of an ensemble model approach allowed the system to overcome the limitations of individual models, ensuring superior stability and achieving higher precision. Furthermore, tests on various real-world biological hypotheses proved the practical utility and reliability of the framework. This study establishes a foundation for precise literature analysis that accounts for biological complexity, serving as a powerful tool to accelerate evidence-based biomedical discoveries.

## 2 Methods

### 2.1 Hypothesis-Driven Evidence Classification Framework

To systematically evaluate scientific evidence, we developed a hypothesis-driven evidence classification framework based on large language models (LLMs). Given a predefined hypothesis and the abstract of a retrieved article, the model determines whether the abstract supports, refutes, or is neutral with respect to the hypothesis.

The target literature set was constructed using a high-recall retrieval strategy. We initially employed keyword-based PubMed queries to maximize sensitivity. However, such queries are limited in handling synonymous or variably expressed biomedical entities. To address this limitation, we incorporated PubTator3-based retrieval, which leverages normalized biomedical entities.[4] This enables the identification of articles that refer to the same concept using different names or synonyms, thereby improving retrieval coverage while preserving the recall-oriented design.

For evidence assessment, only article abstracts were used. Each abstract was classified into one of three categories: *support*, when the abstract provides evidence consistent with the hypothesis; *refute*, when the abstract demonstrates an explicitly opposite directional effect; and *neutral*, when the abstract does not address the hypothesis, provides insufficient evidence, or reports no significant difference. Importantly, null findings were treated as *neutral* rather than *refute*.

As illustrated in Fig. 1, the overall framework integrates high-recall literature retrieval with hypothesis-driven classification and structured output generation.

**Figure 1:**
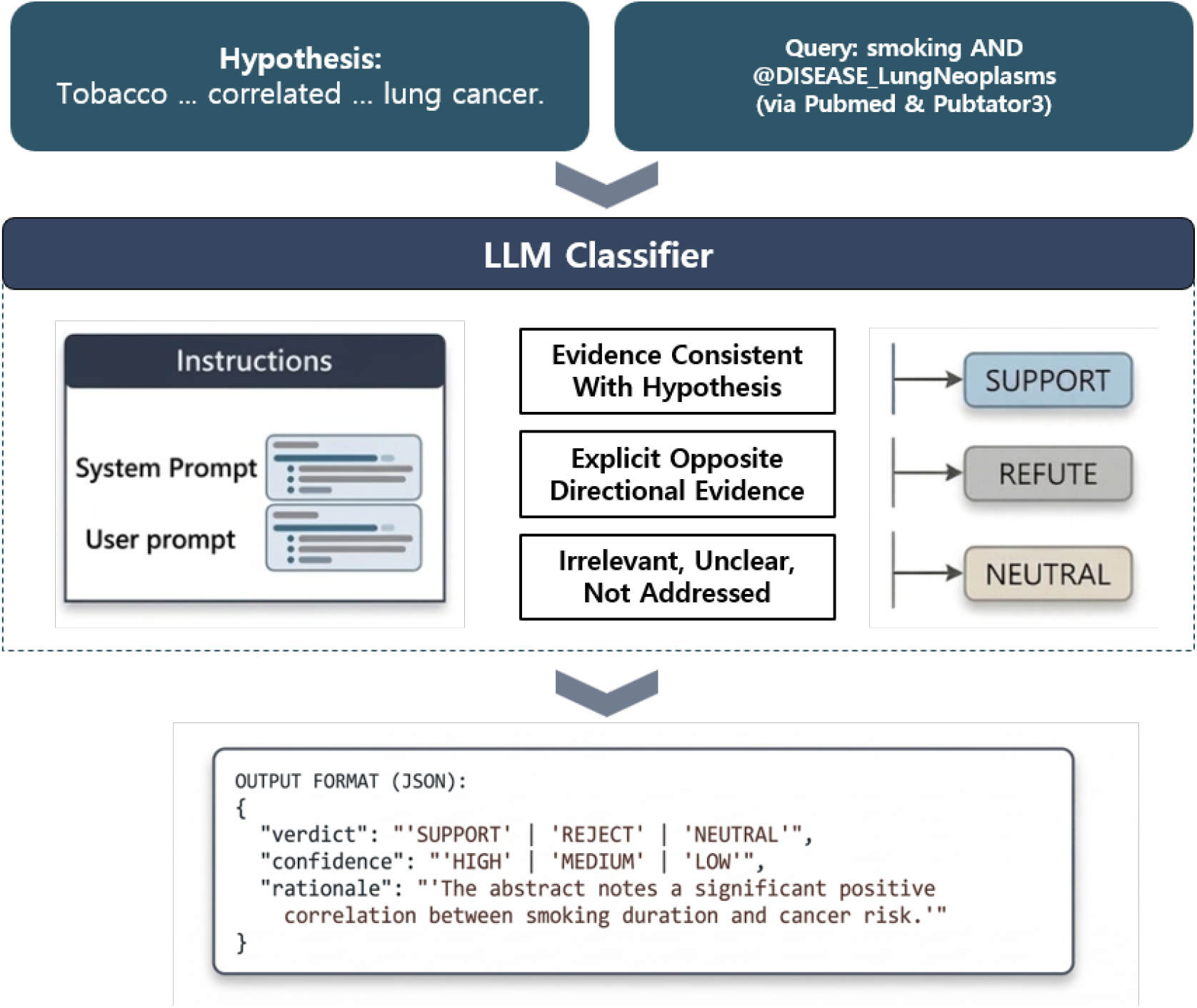
Overview of the hypothesis-driven evidence classification framework.

**Figure 2:**
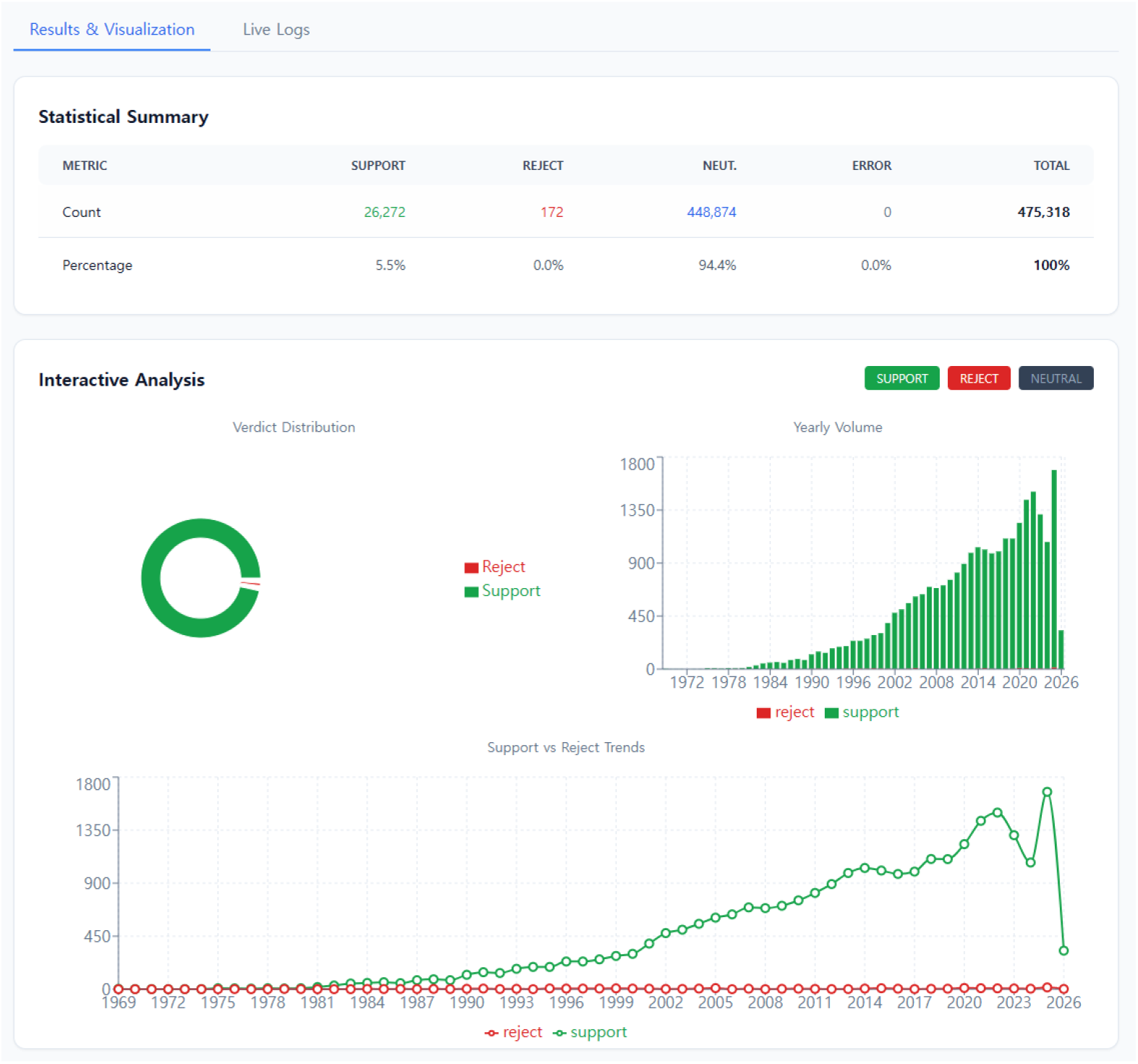
BELIEVE: Web based framework for Bio-medical literature evidence exploration. Example of hypothesis “Patients with Type 2 Diabetes Mellitus exhibit insulin resistance.”

The classification was performed using a structured prompt that enforces consistent evaluation criteria and output formatting. The prompt defines the three evidence categories, explicitly treats null findings as neutral, and requires the model to generate a confidence score along with a brief rationale in structured JSON format. The complete prompt template is provided in the Supplementary Information.

### 2.2 BELIEVE: Web-Based Framework

To operationalize the proposed framework, we developed **BELIEVE (Bio-medical Literature Evidence Exploration)**, a web-based platform available at believe.kaist.ac.kr. BELIEVE implements the hypothesis-driven evidence classification framework described above, providing an end-to-end system for large-scale biomedical literature analysis.

The platform supports flexible dataset construction by integrating both PubMed keyword-based queries and PubTator3-based entity-normalized retrieval, enabling users to efficiently collect and manage high-recall literature sets. Users can define and manage hypotheses, configure model parameters, and execute large-scale evidence classification tasks directly through the interface.

In addition, BELIEVE is supported by a scalable backend architecture designed to handle high-volume traffic and concurrent processing. The system includes modules for model configuration, task scheduling, and structured result storage, ensuring efficient and reproducible analysis.

Through this integrated environment, BELIEVE facilitates systematic exploration, evaluation, and interpretation of biomedical evidence at scale.

## 3 Results

### 3.1 Benchmarking Reasoning Performance on BioNLI

To evaluate the Natural Language Inference (NLI) capabilities of various Large Language Models (LLMs), we utilized the BioNLI dataset [12]. The dataset architecture comprises “true” labels, where the hypothesis is a direct consequence of a matched abstract, and “false” labels, where the hypothesis is adversarial. These adversarial hypotheses were constructed via rule-based perturbations, including *Verb Negation* (VNeg; consisting of posToNeg and negToPos labels) and *Lexical Polarity Reversal* (LPR). Additionally, we generated “neutral” data by randomly shuffling hypotheses and abstracts.

For benchmarking, we randomly sampled 400 data points, consisting of 133 positive, 133 negative (distributed across all three adversarial labels), and 134 neutral labels. Models were accessed via the OpenRouter platform API and instructed to classify whether a given abstract “supports,” “refute,” or is “neutral” toward a specific hypothesis, with outputs constrained to a structured JSON format. Performance was quantified using accuracy, macro-precision, macro-recall, and macro-*F*_1_ scores.[13]

To maximize reliability and mitigate hallucinations, we implemented an ensemble approach using majority voting.[14] We simulated ensembles of varying sizes (*n* = {3, 5, 7, …, 23}) and selected the configuration with the highest accuracy for subsequent literature validation tasks.

We selected state-of-the-art LLM models based on the LiveBench leaderboard across three tasks related to the natural language inference (NLI) problem: reasoning, language, and instruction following (IF).[15] We used the version of the LiveBench leaderboard dated December 23, 2025, and selected the top 23 models out of a total of 48 models (approximately 50%) based on the average of the three task scores. The selected models, along with their specific configurations, are listed in Supplementary Table S3.

#### Benchmarking Capabilities on Individual LLMs

The Natural Language Inference (NLI) capability of the selected models was rigorously evaluated using the **BioNLI dataset** [12]. This dataset architecture is specifically designed to assess whether models can distinguish valid biological claims from subtle semantic inversions and unrelated distractors.

As shown in Figure 3, all 23 evaluated models achieved an accuracy exceeding 0.85 (Supplementary Table S6). The top-performing single model, *gemini-3-pro-preview-11-2025-high*, achieved an accuracy of 0.945, with corresponding *F*_1_-score of 0.9445.

**Figure 3:**
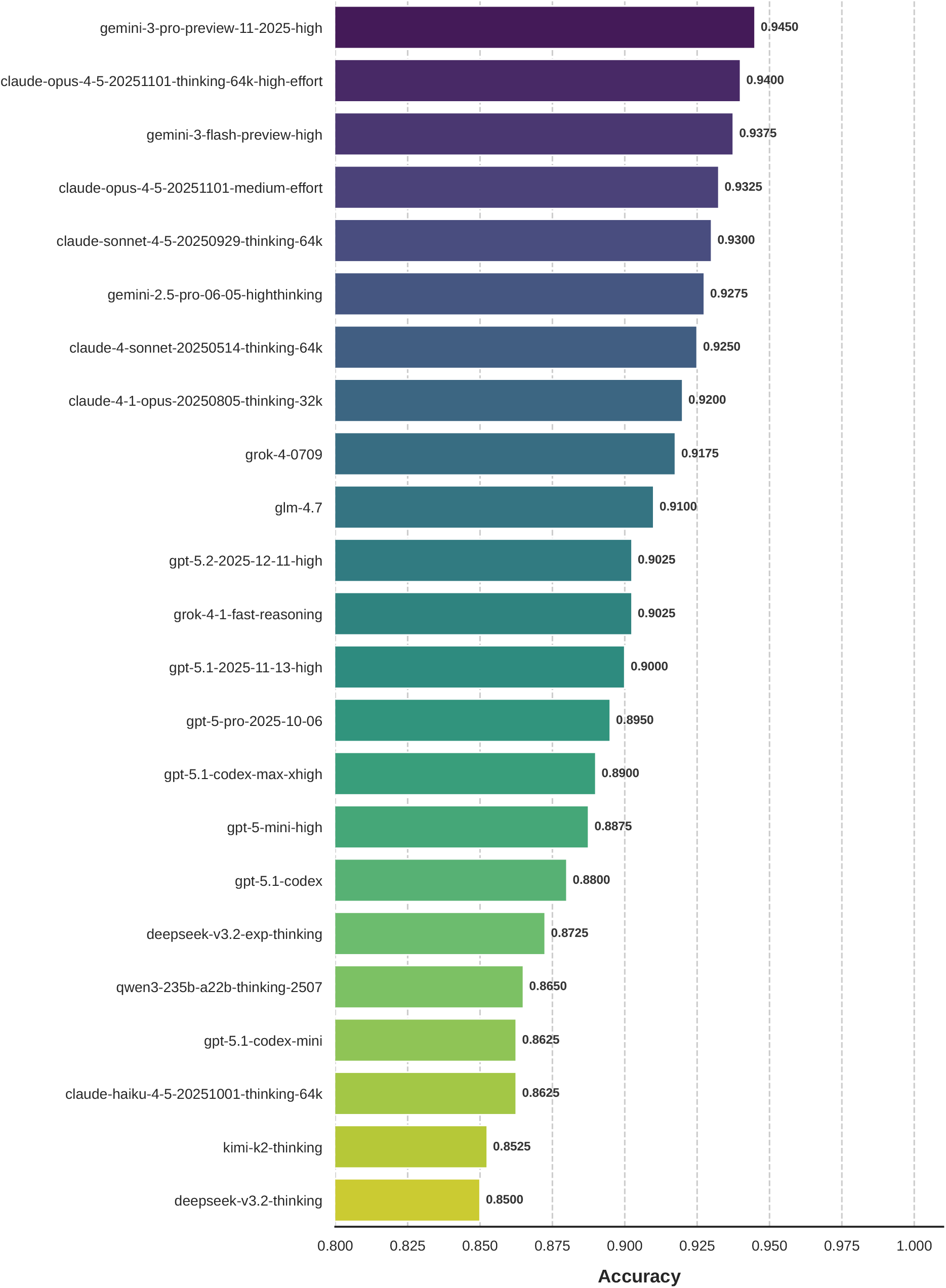
Accuracy comparison of 23 individual models on the BioNLI benchmark.

To further investigate the relationship between general LLM capabilities and BioNLI performance, we computed the Spearman correlation between LiveBench scores (global, reasoning, language, and instruction following) and BioNLI evaluation metrics.[16]

As shown in Figure 4, BioNLI performance exhibits a strong correlation with language capability (Spearman *ρ≈* 0.70), while showing minimal correlation with reasoning and instruction-following performance (*ρ≈* 0.17–0.18). Notably, global benchmark scores also demonstrate weak correlation with BioNLI outcomes, indicating that overall leaderboard rankings are not predictive of domain-specific inference performance.

**Figure 4:**
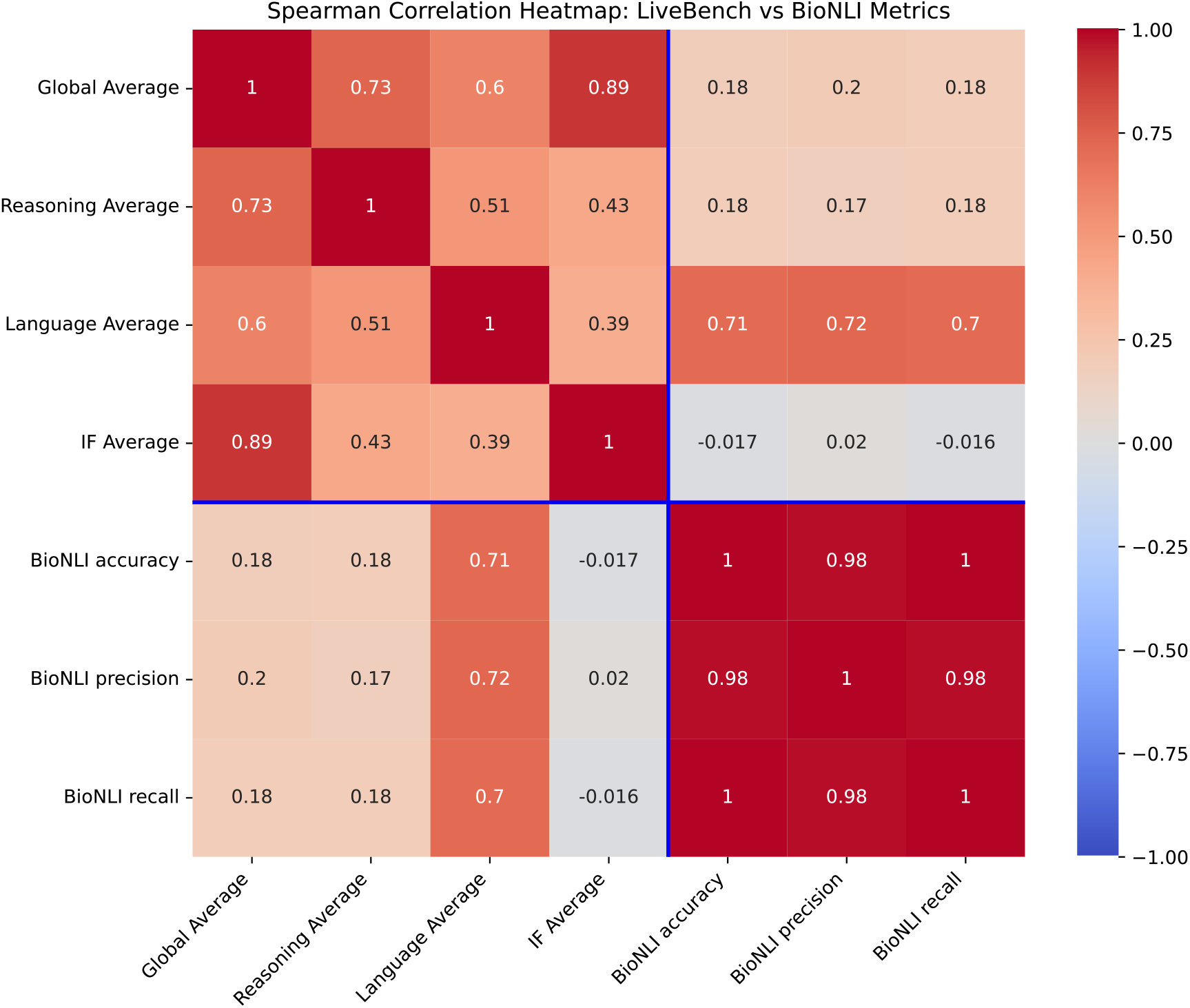
Spearman correlation heatmap between LiveBench scores and BioNLI performance metrics.

These results suggest that BioNLI primarily relies on semantic understanding and linguistic alignment rather than formal reasoning ability. This highlights a fundamental mismatch between general-purpose LLM benchmarks and biologically grounded inference tasks.

#### Ensemble Strategy and Reliability Enhancement

To mitigate model-specific biases, we implemented an ensemble approach based on **majority voting** [14]. As demonstrated in Figure 5(a), increasing the ensemble size significantly stabilized the performance.

**Figure 5:**
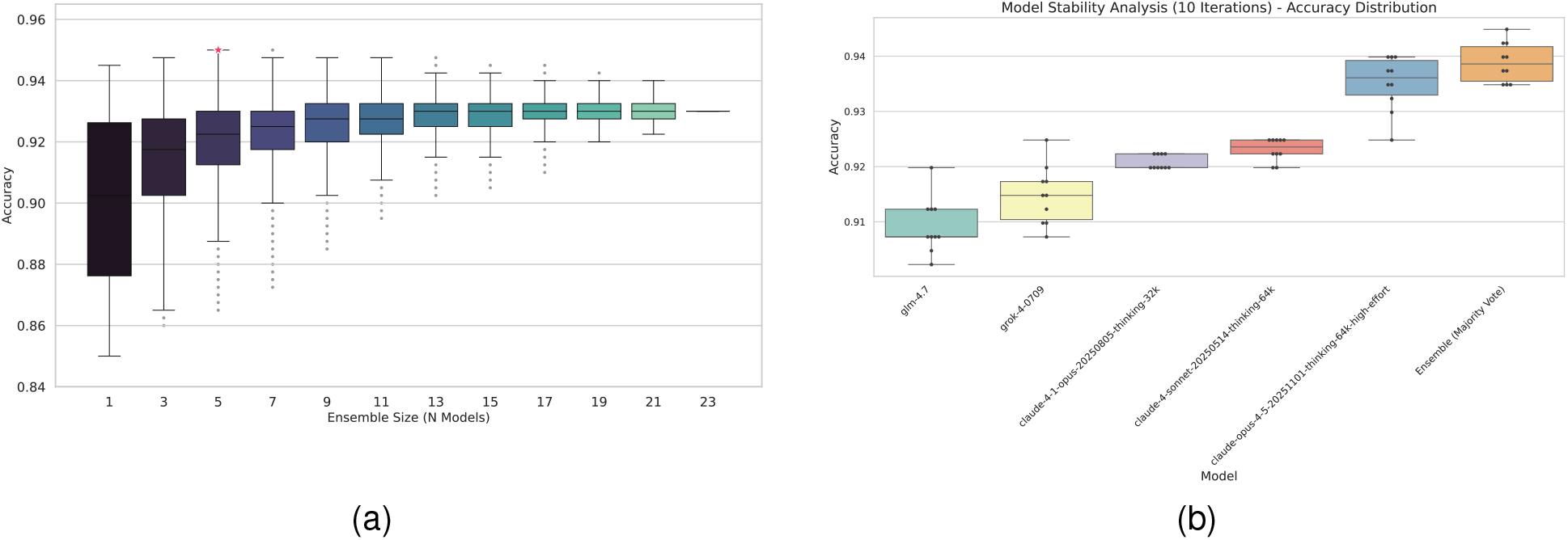
Ensemble performance and stability analysis. **(a)** Reduction in performance variance with increasing ensemble size. **(b)** Stability analysis over 10 iterations for ensemble configurations.

The optimal ensemble was identified as a 5-model configuration (including *claude-4-1-opus* and *grok-4*). This configuration outperformed the best-performing single model across all metrics, confirming that model diversity is more beneficial for alleviating natural variabilities (Figure 5(b)).

#### Consensus Analysis and Replicability

To quantify the strength of agreement, we calculated Fleiss’s kappa (*κ*), yielding a value of **0.9084** [17]. This indicates strong inter-model agreement. While individual models exhibited variable outputs, the ensemble approach demonstrated substantially lower variance, establishing a stable “literature reviewer.”

### 3.2 Validation on Well-Established Biological Hypotheses

To rigorously evaluate the reliability of the proposed framework, we conducted validation experiments on a set of well-established biological hypotheses with known directional relationships. For each hypothesis, we additionally constructed adversarial variants with reversed or negated statements, enabling controlled assessment of whether the model can correctly distinguish supporting and contradicting evidence.

All experiments were performed using the *GLM-4*.*7-FP8* model in combination with PubTator3-based retrieval.[18, 4]

Example hypothesis pair

*Patients with Type 2 Diabetes Mellitus exhibit insulin resistance*.

*Patients with Type 2 Diabetes Mellitus do not exhibit insulin resistance*.

To perform this validation, each hypothesis was converted into a structured literature query using PubTator 3.0, which normalizes biomedical entities and enables precise retrieval from PubMed.[4] Retrieved abstracts were then processed using the LLM-based classification framework to determine whether they support, refute, or are neutral with respect to the hypothesis.

We quantified the results using two key metrics:

- **Relevancy Score** = 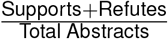: Represents the proportion of abstracts that are relevant to the hypothesis.
- **Alignment Score** = 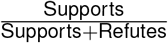: Indicates the degree of scientific consensus supporting the hypothesis.

As shown in Table 1, the framework consistently recovered the correct directional relationships across diverse biological domains. The reported results were obtained from a random sample of 9,000 instances drawn from the full query set. For true hypotheses, the model produced a high proportion of *support* classifications, resulting in near-perfect alignment scores (e.g., 1.0000, 0.9981, 0.9832, 0.9805, 0.9704, and 0.9444). In contrast, adversarial or negated hypotheses were predominantly classified as *refute*, yielding alignment scores close to zero (e.g., 0.0000, 0.0097, 0.0440, 0.0051, 0.0767, and 0.0769).

**Table 1:** Validation on established biological hypotheses and their adversarial variants using the *GLM-4*.*7-FP8* model.

| Hypothesis | Type | Supports | Refutes | Neutrals | Relevancy | Alignment |
| --- | --- | --- | --- | --- | --- | --- |
| T2DM → insulin resistance | True | 1,075 | 0 | 7,923 | 0.1195 | 1.0000 |
| T2DM → insulin resistance | Adversarial | 0 | 1,748 | 7,249 | 0.1943 | 0.0000 |
| Tobacco → lung cancer | True | 522 | 1 | 8,469 | 0.0582 | 0.9981 |
| Tobacco → lung cancer | Adversarial | 7 | 714 | 8,277 | 0.0801 | 0.0097 |
| Metformin → tumor proliferation | True | 644 | 11 | 8,339 | 0.0728 | 0.9832 |
| Metformin → tumor proliferation | Adversarial | 34 | 739 | 8,221 | 0.0859 | 0.0440 |
| Vancomycin → bactericidal activity | True | 301 | 6 | 8,693 | 0.0341 | 0.9805 |
| Vancomycin → bactericidal activity | Adversarial | 3 | 585 | 8,410 | 0.0653 | 0.0051 |
| α-synuclein → neurodegeneration | True | 1,015 | 31 | 7,954 | 0.1162 | 0.9704 |
| α-synuclein → neurodegeneration | Adversarial | 90 | 1,083 | 7,827 | 0.1303 | 0.0767 |
| Amyloid-β → Alzheimer's progression | True | 238 | 14 | 8,744 | 0.0280 | 0.9444 |
| Amyloid-β → Alzheimer's progression | Adversarial | 52 | 624 | 8,320 | 0.0751 | 0.0769 |

**Table 2:** LLM-based evaluation of biological hypotheses using a restricted literature set.

| Hypothesis | # Abstracts | Supports | Refutes | Neutrals | Relevancy | Alignment |
| --- | --- | --- | --- | --- | --- | --- |
| H1 | 140,629 | 1,075 | 0 | 7,923 | 0.1195 | 1.0000 |
| H2 | 140,629 | 0 | 1,748 | 7,249 | 0.1943 | 0.0000 |
| H3 | 78,665 | 522 | 1 | 8,469 | 0.0582 | 0.9981 |
| H4 | 78,665 | 7 | 714 | 8,277 | 0.0801 | 0.0097 |
| H5 | 34,576 | 644 | 11 | 8,339 | 0.0728 | 0.9832 |
| H6 | 34,576 | 34 | 739 | 8,221 | 0.0859 | 0.0440 |
| H7 | 21,714 | 301 | 6 | 8,693 | 0.0341 | 0.9805 |
| H8 | 21,714 | 3 | 585 | 8,410 | 0.0653 | 0.0051 |
| H9 | 9,496 | 1,015 | 31 | 7,954 | 0.1162 | 0.9704 |
| H10 | 9,496 | 90 | 1,083 | 7,827 | 0.1303 | 0.0767 |
| H11 | 17,637 | 238 | 14 | 8,744 | 0.0280 | 0.9444 |
| H12 | 17,637 | 52 | 624 | 8,320 | 0.0751 | 0.0769 |

**Table 3:**
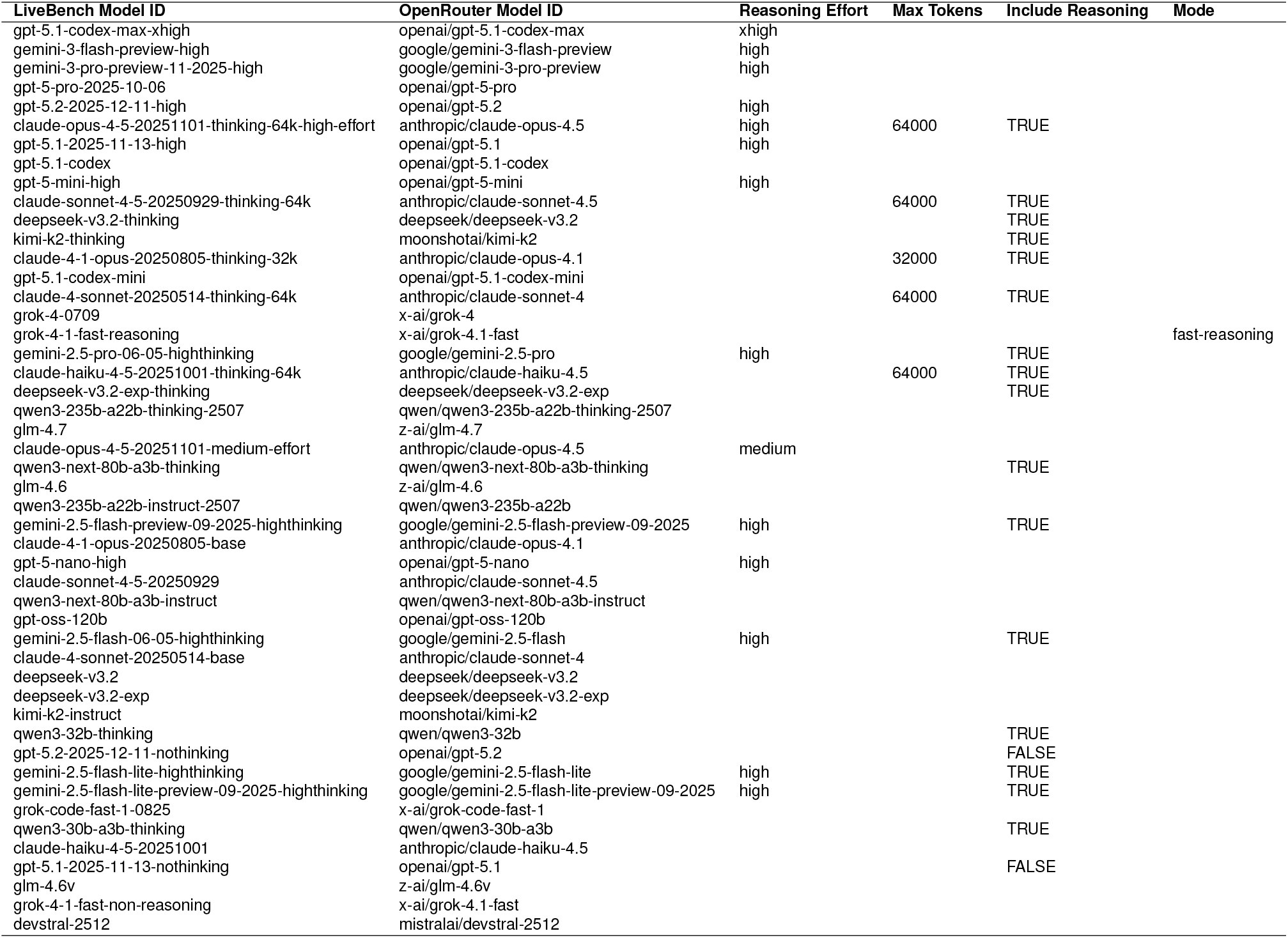
Supplementary Table S3. Selected LLMs and their OpenRouter configurations, including reasoning effort, maximum token limits, reasoning inclusion, and inference modes.

This pattern was consistently observed across multiple biological domains, including metabolic disorders, epidemiological associations, pharmacological effects, antimicrobial mechanisms, and neurodegenerative disease pathways.

These results demonstrate that the framework not only retrieves relevant evidence but also accurately captures the directionality of biological relationships. The clear separation between true and adversarial hypotheses further indicates robustness against misleading or contradictory formulations, supporting the validity of the proposed approach for systematic evidence analysis.

## 4 Discussion

This study presents a hypothesis-driven framework for automated biomedical literature analysis that evaluates evidence at the level of individual publications. Unlike conventional RAG-based approaches, which often lose critical biological context and favor generalized conclusions, our method preserves contextual integrity by requiring abstract-level, instance-wise evaluation. The results demonstrate that the framework can reliably identify both supporting and contradictory evidence, with strong performance on the BioNLI task and robust validation across established biological hypotheses. The ensemble approach further improved stability and reduced model-specific variability, achieving high inter-model agreement and consistent classification outcomes. Importantly, these findings suggest a practical strategy for real-world applications: when the optimal model for a given task is uncertain, leveraging a set of state-of-the-art models in an ensemble configuration can provide more reliable and robust performance than relying on a single model.

Notably, our correlation analysis reveals that BioNLI performance is strongly associated with language understanding rather than reasoning capability, challenging the assumption that general reasoning benchmarks directly translate to scientific inference performance. In addition, the performance gap across models was not substantial, suggesting that models with strong language capabilities can be effectively utilized without strict dependence on a specific architecture or ranking. This finding highlights a fundamental mismatch between current LLM evaluation paradigms and the requirements of biomedical evidence synthesis. Beyond classification, the proposed framework provides a foundation for systematically analyzing both supporting and contradictory evidence, enabling the identification of the specific biological contexts in which a given hypothesis holds or fails. Such analysis can be further extended to uncover the conditions under which broader biological phenomena emerge, offering a scalable approach to context-aware knowledge synthesis in complex biological systems.

## 5 Data availability

All data supporting the findings of this study are available in the BELIEVE GitHub repository at

https://github.com/bisl-hub/believe.

## 6 Code availability

The source code for the proposed framework, including the BELIEVE platform, is publicly available at https://github.com/bisl-hub/believe.

## 7 Acknowledgements

This research was supported by the Bio&Medical Technology Development Program of the National Research Foundation (NRF) funded by the Korean government (MSIT) (No. RS-2025-16063391)

## Supplementary Information

### 8 Supplementary Information

#### S1. LLM Prompt for Evidence Classification

The full prompt used for hypothesis-driven evidence classification is provided below.

~~~
You are a biomedical literature reviewer. Classify whether the given ABSTRACT supports, contradicts, or is neutral toward the HYPOTHESIS. Base your reasoning strictly on the ABSTRACT.
EVALUATION CATEGORIES:
SUPPORT: The abstract provides evidence that the hypothesis is correct REJECT: The abstract explicitly demonstrates the opposite direction
NEUTRAL: The abstract shows no significant difference, unclear results, or does not address any phenomenon related to the hypothesis
CRITICAL CLARIFICATION:
No significant difference or no difference shown = NEUTRAL (null finding, not refutation)
Abstract does not mention the phenomenon = NEUTRAL (no evidence to evaluate) Only classify as REJECT when an explicit opposite directional effect is shown
CONFIDENCE LEVELS:
HIGH: Clear evidence directly addresses the hypothesis with explicit directional findings
MEDIUM: Relevant findings present but with some ambiguity or indirect relevance LOW: Minimal evidence, borderline relevance, or high interpretive uncertainty
OUTPUT FORMAT (JSON):
{“verdict”: “[SUPPORT | REJECT | NEUTRAL ]”,
“confidence”: “[HIGH | MEDIUM | LOW ]”,
“rationale”: “[justification referencing the abstract, about 2–3 sentences ]” }
~~~

#### S2. Evaluated Biological Hypotheses

##### Hypotheses and PubTator Queries

- **H1**: Patients with Type 2 Diabetes Mellitus exhibit insulin resistance. *Query:* @DISEASE_Diabetes_Mellitus_Type_2 AND @DISEASE_Insulin_Resistance
- **H2**: Patients with Type 2 Diabetes Mellitus do not exhibit insulin resistance. *Query:* @DISEASE_Diabetes_Mellitus_Type_2 AND @DISEASE_Insulin_Resistance
- **H3**: Tobacco smoking is positively correlated with an increased incidence of lung cancer. *Query:* smoking AND @DISEASE_Lung_Neoplasms
- **H4**: Tobacco smoking is negatively correlated with an increased incidence of lung cancer. *Query:* smoking AND @DISEASE_Lung_Neoplasms
- **H5**: Metformin demonstrates anti-proliferative effects on tumor cells. *Query:* @CHEMICAL_Metformin AND @DISEASE_Neoplasms
- **H6**: Metformin fails to demonstrate anti-proliferative effects on tumor cells. *Query:* @CHEMICAL_Metformin AND @DISEASE_Neoplasms
- **H7**: Vancomycin exerts bactericidal activity via peptidoglycan biosynthesis inhibition. *Query:* @CHEMICAL_Vancomycin AND @DISEASE_Bacterial_Infections
- **H8**: Vancomycin does not exert bactericidal activity and is ineffective in Gram-positive infections. *Query:* @CHEMICAL_Vancomycin AND @DISEASE_Bacterial_Infections
- **H9**: Alpha-synuclein aggregation mediates neurodegeneration in Parkinson’s disease. *Query:* synuclein AND @DISEASE_Parkinson_Disease AND substantia nigra AND @CHEM-ICAL_Dopamine
- **H10**: Alpha-synuclein aggregation does not mediate neurodegeneration in Parkinson’s disease. *Query:* synuclein AND @DISEASE_Parkinson_Disease AND substantia nigra AND @CHEM-ICAL_Dopamine
- **H11**: Amyloid-beta deposition contributes to Alzheimer’s disease progression. *Query:* amyloid AND @DISEASE_Alzheimer_Disease AND frontal cortex
- **H12**: Amyloid-beta deposition does not contribute to Alzheimer’s disease progression. *Query:* amyloid AND @DISEASE_Alzheimer_Disease AND frontal cortex

##### LLM Evaluation Results

#### Supplementary Table S3. Selected LLMs and OpenRouter configurations

We provide the list of selected LLMs from the LiveBench leaderboard along with their corresponding OpenRouter model identifiers and inference configurations, including reasoning effort, maximum token limits, reasoning inclusion, and inference modes.

